# Effect of Platelet-activating factor on barrier function of ARPE-19 cells

**DOI:** 10.1101/398131

**Authors:** Fan Zhang, Lei Liu, Han Zhang, Zhe-Li Liu

**Author notes:** **Corresponding author:** Han Zhang, M.D., Ph.D., Department of Ophthalmology, the First Affiliated Hospital of China Medical University, 155 Nanjing North St, Heping District, Shenyang 110001, Liaoning, People’s Republic of China.

## Abstract

**PURPOSE:** To examine the effects of platelet-activating factor (PAF) on tight junction permeability in cultured retinal pigment epithelial (RPE) cells.

**METHODS:** A human RPE cell line (ARPE-19) cultured on microporous filter supports was used. PAF and WEB2086, which is a specific PAF-receptor (PAF-R) antagonist, were added to the culture medium. RPE monolayer permeability was measured using transepithelial electrical resistance (TER) and sodium fluorescein flux. The expression of the tight junction protein zonula occludens (ZO)-1 was assessed using immunohistochemistry. We also measured the vascular endothelial growth factor (VEGF) level in cultures treated with PAF, and RPE monolayer permeability was measured again in the presence of neutralizing antibodies to VEGF.

**RESULTS:** PAF significantly decreased the TER of the RPE monolayer and enhanced sodium fluorescein flux. ZO-1 expression was downregulated in PAF-supplemented medium. These effects were abolished with PAF-R blockage. PAF stimulation increased VEGF expression in RPE cells, and neutralization of VEGF with antibodies caused partial recovery of barrier properties.

**CONCLUSIONS:** The tight junctions of ARPE-19 cells are altered by PAF, and these effects are partly mediated by the upregulation of VEGF expression in these cells. Our results contribute to growing evidence supporting the role of PAF in choroidal neovascularization, and our findings suggest that PAF is a novel therapeutic target for increased permeability of the RPE monolayer.

## 1. Introduction

The retinal pigment epithelium (RPE) is a single monolayer of epithelial cells at the back of the eye that forms the outer blood-retinal barrier (BRB). It is fundamentally important for the health and integrity of the distal retina. The RPE plays important roles in the maintenance of the visual cycle and phagocytosis of the photoreceptor outer segment. Additionally, it is the main transport pathway for the flow of nutrients, metabolic waste products, ions, and fluid between the distal retina and the choriocapillaris[1]. Apical tight junctions that join adjacent RPE structures maintain continuity of the barrier between cells, and they are critical for maintaining normal polarized functions of the RPE monolayer[2]. Barrier breakdown can occur in ocular trauma and many other ocular diseases, such as proliferative vitreoretinopathy, diabetic retinopathy, and neovascular age-related macular degeneration (AMD), leading to increased permeability of serum components. In neovascular AMD, choroidal neovascularization (CNV) initially occurs under Bruch’s membrane and the RPE monolayer and then advances to the subretinal space, leading to subretinal hemorrhage, exudative lesions, serous retinal detachment, and ultimately disciform scarring. It is believed that the equilibrium of polarized secretion of antiangiogenic and proangiogenic factors and then maintenance of the resulting chemotactic gradient owing to the barrier properties of the RPE monolayer play critical roles in the prevention of CNV development in the neurosensory retina[3]. Dysfunction of the outer BRB may lead to photoreceptor degeneration and blindness[4]. However, the mechanisms implicated in RPE barrier regulation have been poorly explored.

The platelet-activating factor (PAF, 1-O-alkyl-2-acetyl-sn-glycero-3-phosphocholine) is considered to be the first bioactive lipid ever identified and is a potent proinflammatory mediator involved in cellular activation, intracellular signaling, apoptosis, diverse inflammatory reactions, and angiogenesis[5-9]. Its biological actions are mediated through the activation of a G protein-coupled PAF receptor (PAF-R)[10]. A recent study showed that a PAF-R is present in RPE cells and choroidal endothelial cells and that PAF increases the production of vascular endothelial growth factor (VEGF) in RPE cells[11]. In addition, our previous data indicated that local expression of PAF-R in the subretinal space is upregulated during experimental CNV development and that administration of a PAF-R antagonist potently reduced CNV lesion size[12]. These findings suggested that PAF might play a role in the pathogenesis of CNV and neovascular AMD. Previous studies have shown that PAF challenge induced endothelial actin cytoskeletal rearrangement and marked endothelial barrier dysfunction[13]; however, the role of PAF in RPE barrier regulation is unclear.

The present study aimed to examine the effects of PAF on the barrier function of cultured RPE cells. Our findings will contribute to growing evidence on the role of PAF in choroidal neovascularization and possibly suggest PAF as a novel therapeutic target for increased permeability of RPE cells.

## 2. Materials and Methods

### 2.1. RPE cell culture

ARPE-19 cells (American Type Culture Collections, Manassas, VA) were cultured as described previously[14]. Briefly, ARPE-19 cells were cultured in Dulbecco’s modified Eagle’s medium (Sigma Chemical Co., St. Louis, MO) containing 10% heat-inactivated fetal bovine serum (FBS; Gibco, Carlsbad, CA) supplemented with 100 U/mL penicillin, 100 mg/mL streptomycin, 1% L-glutamine, and 0.1 mM nonessential amino acids in a humidified incubator with 5% CO_2_ at 37°C. After cell attachment, the medium was modified to contain only 1% FBS and was changed every 2 to 3 days.

### 2.2. Measurement of transepithelial electrical resistance (TER)

To establish cell monolayers, ARPE-19 cells were grown on microporous filter membranes (0.4-μm pore size; Corning Inc., Corning, NY), and the membranes were placed in 24-well culture plates. The TER was measured using an epithelial voltmeter (EVOM; World Precision Instruments, Sarasota, FL), according to the manufacturer’s instructions. The TER values (Ω/cm^2^) were determined by subtracting the resistance of the filter alone (background) from the values obtained with the filters and RPE cells. Measurements were performed every 3 days in the first 16 days and every day thereafter. After TER stabilization, the ARPE-19 monolayer was incubated with an exogenous stable derivative of PAF (carbamyl-PAF [cPAF], 100 nM; Enzo Biochemical, Inc., New York, NY) or cPAF with WEB 2086 (10 μg/ml; Santa Cruz Biotechnology, Santa Cruz, CA). The medium with or without stimulants was changed every other day. Measurements were repeated at least five times for each well, and each experiment was repeated in at least four different wells.

### 2.3. Permeability assay

RPE cell permeability was assessed by measuring the apical-to-basolateral movements of fluorescein isothiocyanate (FITC)-dextran (4 kDa; Sigma Chemical Co.), as previously described[14]. Briefly, 3 weeks after starting culturing, the ARPE-19 monolayer was treated with cPAF (100 nM) or cPAF with WEB 2086 (10 μg/ml) for 6 days. The medium was changed every 2 days, and 1000 μg/ml of FITC-dextran was added to the upper chamber on day 6 following stimulation. Samples (100 μl) were obtained from the upper and lower chambers 24 h after addition of FITC-dextran. The concentration of FITC-dextran in these samples was quantified using spectrophotometry. The diffusion rate was expressed as a percentage and was calculated as follows: (amount of dextran in the lower chamber) × 100 / (amount of dextran in the upper chamber). Each experiment was repeated four times.

### 2.4. Immunohistochemistry

The formation of tight junctions in ARPE-19 cells was examined by staining of ZO-1, which is a tight junction protein critical to barrier integrity. ARPE-19 cells cultured under conditions identical to those in the permeability assay were fixed with 4% paraformaldehyde in phosphate-buffered saline (PBS) for 10 min at room temperature (RT) and rinsed thrice. The ARPE-19 monolayer on filters was permeabilized with 0.2% Triton X-100 in PBS for 20 min and blocked with 10% goat serum at RT for 30 min. Rabbit anti-ZO-1 antibodies (1:100; Invitrogen, Carlsbad, CA) were added, and the mixture was incubated overnight at 4°C. After washing with PBS thrice, the cells were further incubated with FITC-conjugated goat anti-rabbit immunoglobulin (1:200; Invitrogen) for 30 min at RT. The slides were treated with a nuclear stain and were examined under a fluorescent microscope (BZ-9000, Keyence, Japan).

### 2.5. VEGF enzyme-linked immunosorbent assay (ELISA)

The modulation of VEGF production by exogenous PAF was assessed using an ELISA kit (R&D Systems, Minneapolis, MN). RPE cells were incubated in the presence of cPAF (100 nM) or cPAF with WEB 2086 (10 μg/ml) for 24 h. Cells were harvested following trypsinization. The VEGF concentration under various conditions was determined according to the manufacturer’s protocol. Each assay was repeated thrice.

### 2.6. Neutralization of VEGF

Three weeks after starting culturing, the ARPE-19 monolayer was treated with bevacizumab (0.25 mg/ml, Avastin; Hoffmann La Roche, Basel, Switzerland), which is a recombinant humanized monoclonal antibody against all VEGF isoforms, in the presence of cPAF (100 nM) for 6 days. TER measurement, permeability assay, and ZO-1 staining were performed to determine the role of VEGF in RPE barrier dysfunction induced by PAF.

### 2.7. Statistical analysis

Each result is representative of at least three independent experiments. All values are presented as mean ± SEM. Statistical significance was assessed using Student’s *t*-test (SPSS, Chicago, IL). A *P*-value <0.05 was considered to indicate statistical significance.

## 3. Results

### 3.1. PAF reduces TER and increases ARPE-19 monolayer permeability

ARPE-19 monolayers were established on permeable membrane filters. The TER of the ARPE-19 monolayer increased rapidly during the initial 18 days of culture and reached a plateau within the following 3 days. The TER values were similar to those reported previously for this cell line[14-16]. A mean TER of 44.0 ± 3.5 Ω/cm^2^ was recorded on day 22, where after cPAF was added. When compared with the control group, incubation of ARPE-19 monolayers with cPAF induced a gradual decrease in TER, and a significant effect was noted 4 days after stimulation (p < 0.05). Continuous decreases were also observed 5 and 6 days after stimulation (p < 0.01). The effect of cPAF was inhibited by the special PAF receptor antagonist WEB 2086 (Fig. 1). The ARPE-19 monolayer permeability was further evaluated by measuring the transepithelial diffusion rate of FITC-dextran through the monolayer. Stimulation of the ARPE-19 monolayer with cPAF for 6 days induced a higher FITC-dextran diffusion rate at 24 h when compared to that in the control group (p < 0.05). Additionally, the effect of cPAF was inhibited by WEB 2086 (Fig. 2).

**Figure 1.**
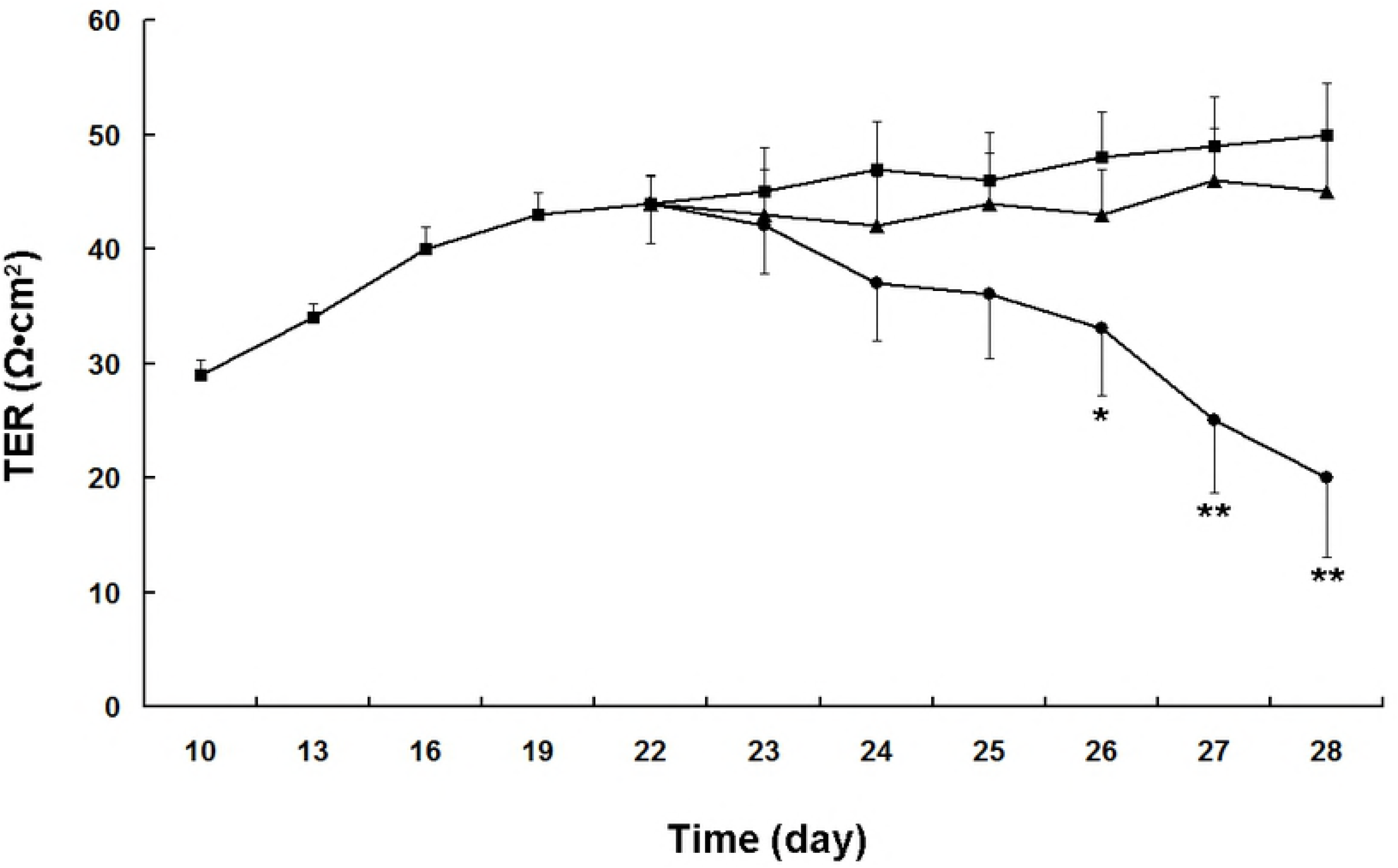
Effect of carbamyl-platelet-activating factor (cPAF) on the transepithelial electrical resistance (TER) of a cultured ARPE-19 monolayer. Monolayers were cultured for 22 days and then cPAF was added. Compared with the control group (▪), incubation of ARPE-19 monolayers with 100 nM of cPAF (•) induced a gradual decrease in TER, and a significant effect was noted 4 days after stimulation (p < 0.05). Continuous decreases were also noted 5 and 6 days after stimulation (p < 0.01). The effect of cPAF was inhibited by the special PAF receptor antagonist WEB 2086 (▴). Data are presented as mean ± SEM. *p < 0.05, **p < 0.01 vs. the control group

**Figure 2.**
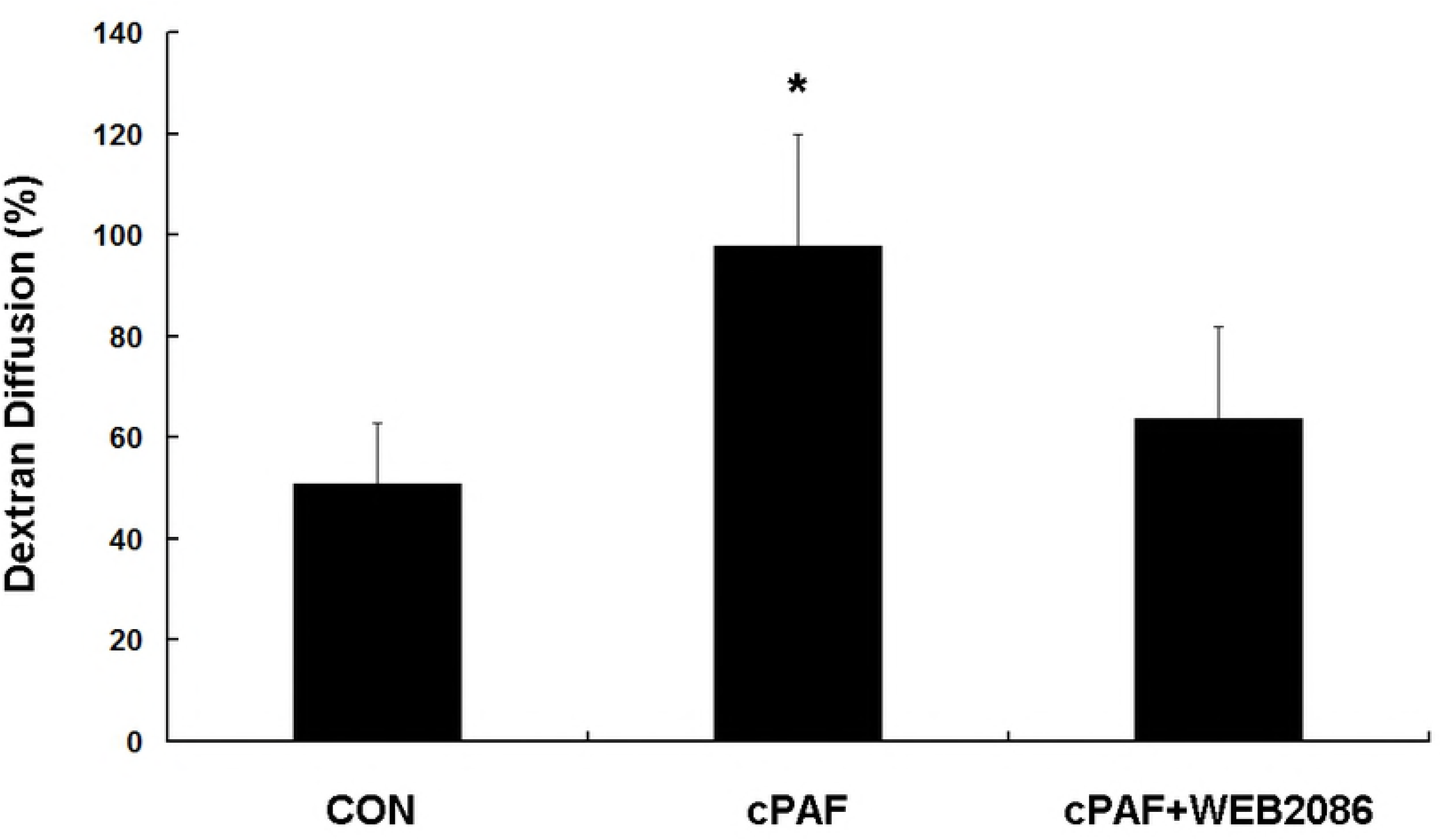
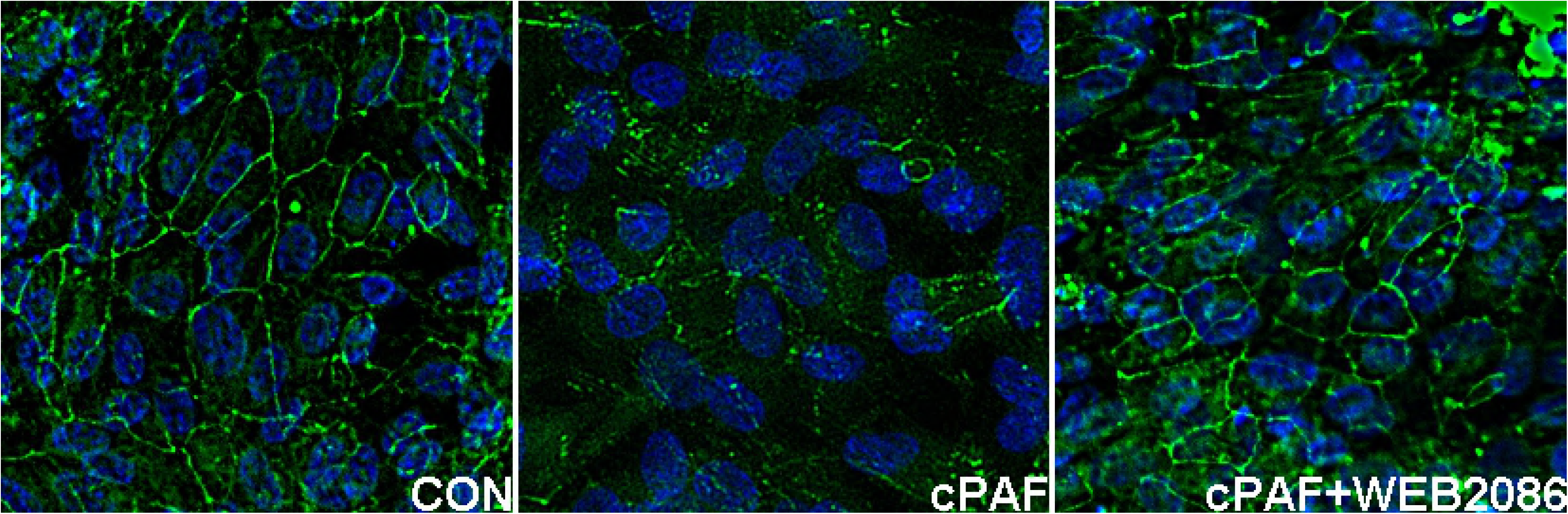
Effect of carbamyl-platelet-activating factor (cPAF) on the transepithelial diffusion rate of FITC-dextran in an ARPE-19 monolayer. Stimulation of an ARPE-19 monolayer with 100 nM of cPAF for 5 days induced a higher FITC-dextran diffusion rate at 24 h when compared with the rate in the control group. The effect of cPAF was inhibited by WEB 2086. A diffusion percentage approaching 100% indicates that the amounts of dextran-FITC in the upper and lower chambers approach the same value. Data are shown as mean ± SEM. *p < 0.05 vs. the control group

### 3.2. PAF causes disruption of ZO-1 distribution in the ARPE-19 monolayer

Barrier function of RPE cells was then evaluated with ZO-1 staining. Immunostaining for ZO-1 in the untreated ARPE-19 monolayer showed continuous labeling in the region of cell-cell contact. Exposure of these cells to cPAF for 6 days caused a markedly disturbed distribution of ZO-1 (Fig. 3).

**Figure 3.**
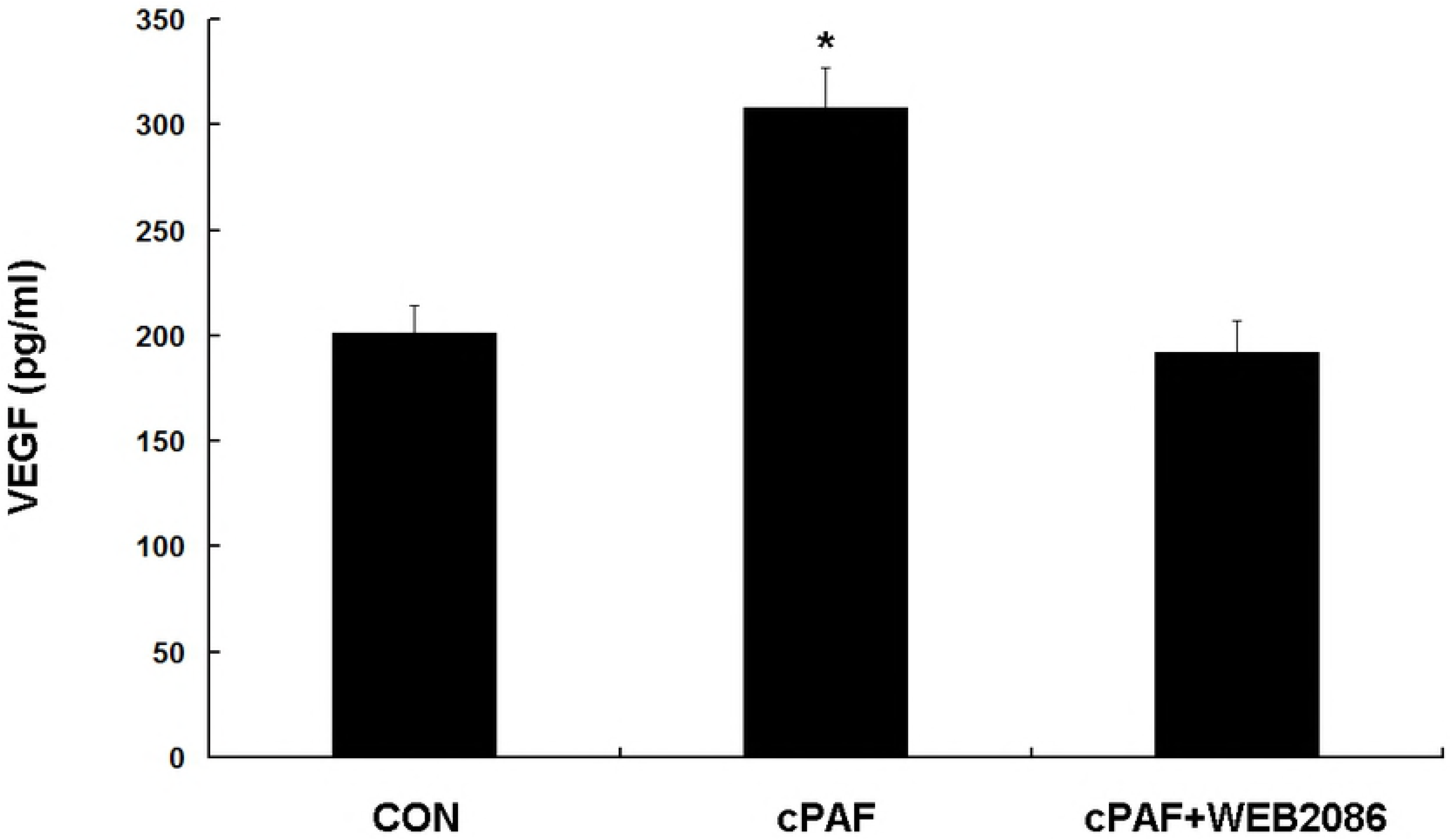
Effect of carbamyl-platelet-activating factor (cPAF) on the distribution of junction proteins in an ARPE-19 monolayer. Cells were incubated with 100 nM of cPAF for 5 days and were then fixed and immunolabeled with ZO-1. Immunostaining for ZO-1 in an untreated ARPE-19 monolayer showed continuous labeling in the region of cell-cell contact. Incubation with cPAF caused marked disruption of ZO-1 staining. The effect of cPAF was inhibited by WEB 2086.

### 3.3. PAF increases VEGF expression in the ARPE-19 monolayer

The VEGF level was significantly higher in RPE cells preincubated with cPAF than in control cells (p < 0.05), and WEB 2086 reduced VEGF expression (Fig. 4).

**Figure 4.**
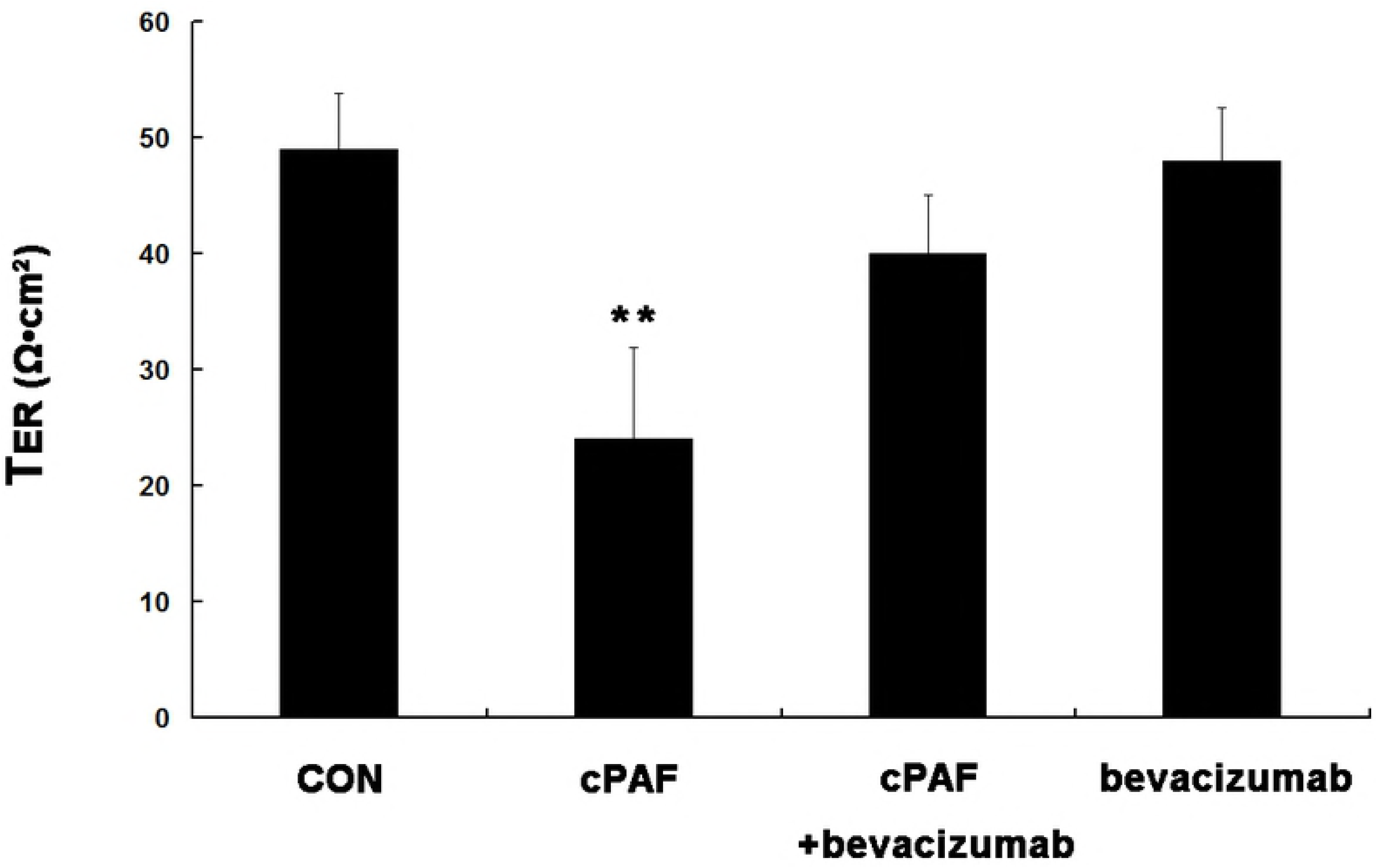
Effect of carbamyl-platelet-activating factor (cPAF) on the vascular endothelial growth factor (VEGF) level in an ARPE-19 monolayer. Established retinal pigment epithelial cell monolayers were incubated with 100 nM of cPAF for 24 h, and then, the modulation of VEGF production by exogenous cPAF was assessed using ELISA. The VEGF level in the presence of cPAF was significantly higher than that in the control group. The effect of cPAF was inhibited by WEB 2086. Data are shown as mean ± SEM. *p < 0.05 vs. the control group

### 3.4. Neutralization of VEGF led to partial recovery of barrier dysfunction induced by PAF

VEGF blockage in the ARPE-19 monolayer significantly prohibited a reduction in TER (p < 0.01) and increase in permeability (p < 0.05) induced by cPAF on day 6 (Fig. 5A, B). Similarly, VEGF blockage attenuated cPAF-induced disruption of ZO-1 staining (Fig. 5C). Treatment with bevacizumab alone did not affect cPAF-induced changes in TER, permeability, and ZO-1 distribution in the ARPE-19 monolayer.

**Figure 5.**
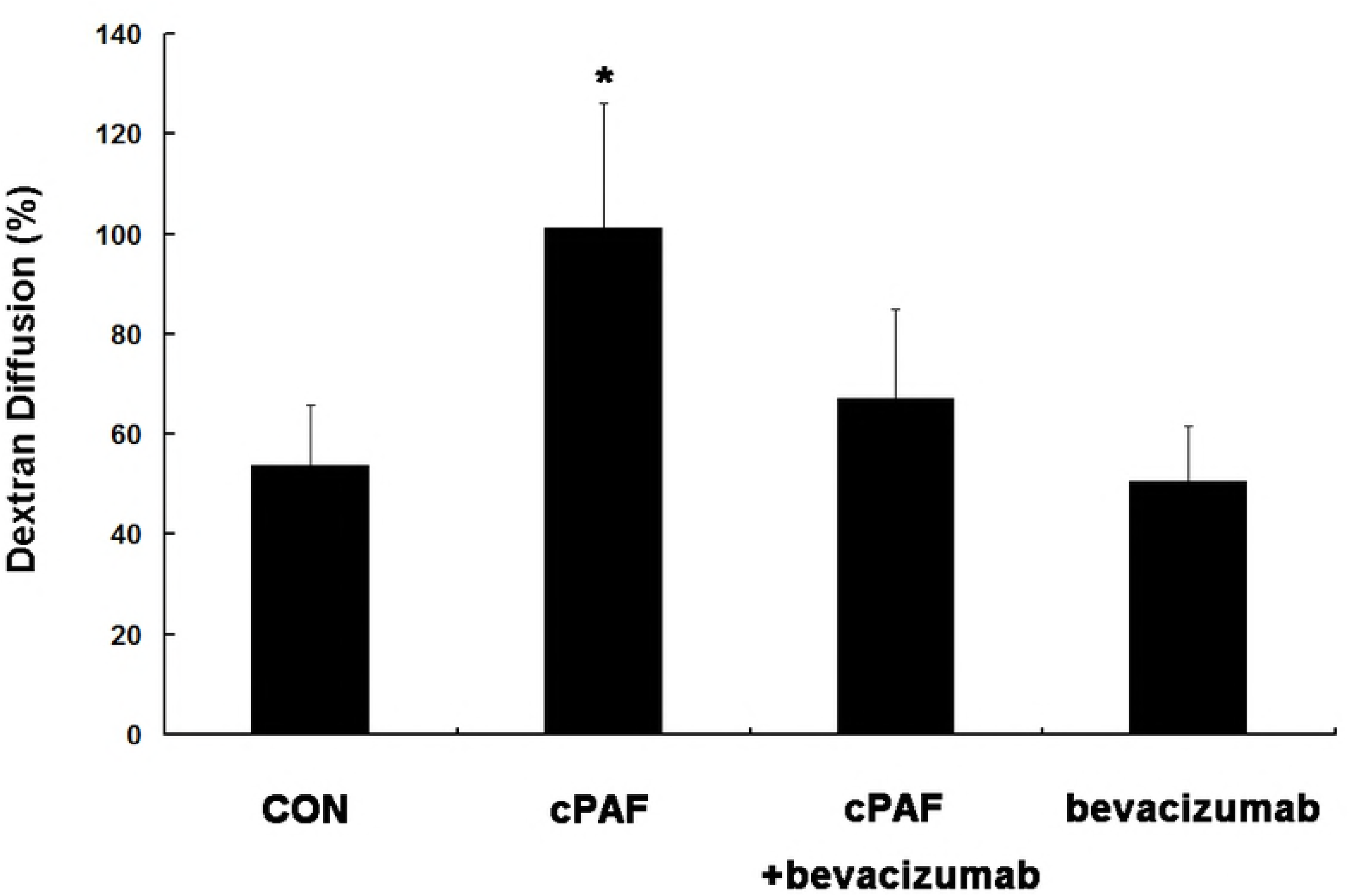
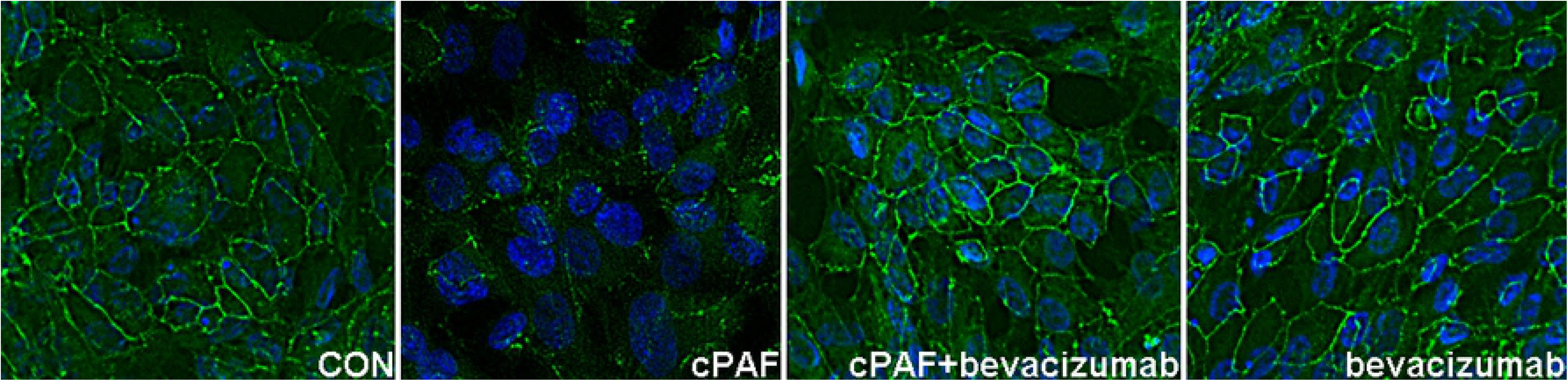
Neutralization of vascular endothelial growth factor (VEGF) with bevacizumab caused the recovery of barrier dysfunction induced by carbamyl-platelet-activating factor (cPAF), indicating that PAF modulation of retinal pigment epithelial (RPE) barrier function is at least partly mediated by VEGF release from RPE cells.

## 4. Discussion

The present study found that PAF significantly decreased the TER of the RPE monolayer and enhanced sodium fluorescein flux. The expression of ZO-1 was downregulated in PAF-supplemented medium. These effects were abolished by PAF-R blockage. PAF stimulation increased the VEGF level in RPE cells, and neutralizing VEGF with antibodies caused partial recovery of barrier properties. To our knowledge, the present study is the first to characterize the relationship between PAF and the barrier function of cultured RPE cells.

The ARPE-19 cell line was used to study the mechanisms responsible for PAF-mediated changes in RPE permeability. ARPE-19 is a human RPE cell line with differentiated properties, and it exhibits morphological polarization when cultured as a monolayer on microporous filter supports[17]. To evaluate the integrity of the RPE barrier, the TER assay, which is a classic method for measuring the resistance to an electrical current across a monolayer[18,19], was used. Under the conditions mentioned above, the tight junctions of the RPE cells had a maximum TER of 50 to 100 Ω/cm^2^[14-16]. In the present study, the TER of the ARPE-19 monolayer gradually increased and reached a plateau of 44 Ω/cm^2^ after 3 weeks of culture. This finding was consistent with the results of previous studies[14-16]. Addition of PAF resulted in a decrease in the TER of the ARPE-19 monolayer. Transepithelial fluxes of FITC-dextran, which is a test to evaluate the structural integrity of cultured epithelium monolayers[14,19,20], was used in the present study. Exposure to PAF for 5 days resulted in a significantly increased diffusion rate as compared to that in unstimulated controls. These results suggest that PAF could significantly compromise the barrier function of the ARPE-19 monolayer and that it had an effect on the structural integrity of the monolayer.

The barrier function of the epithelium depends on intercellular tight junctions involving a number of tight junction proteins. ZO-1 has been shown to be exclusively localized in tight junction strands in various types of epithelial cells and is considered as one of the most important tight junction proteins of the RPE cell layer[13]. Our results showed that the expression of ZO-1 was lower in the presence of PAF on immunohistochemistry. Abnormal distribution of the ZO-1 protein caused by PAF may result in decreased TER and an increased diffusion rate of FITC-dextran. A previous study showed that PAF could induce severe endothelial barrier leakiness by causing the redistribution of ZO-1 and other junctional proteins inside endothelial cells[13]. Similarly, Xu et al. reported that PAF played an important role in mucosal permeability regulation, and the effects were correlated with structural alterations of the F-actin-based cytoskeleton and tight junction proteins, including ZO-1, in intestinal epithelial cells[21].

VEGF has been implicated in the disruption of RPE barrier function. Ablonczy et al. demonstrated that VEGF administration caused a significant drop in TER among ARPE-19 cells[16]. On the other hand, the PAF-R was noted in RPE cells, and PAF increased VEGF production in RPE cells[11]. Considering these findings, we propose that VEGF is a likely candidate associated with PAF-induced RPE barrier dysfunction. We confirmed that PAF increased VEGF expression in the ARPE-19 monolayer and found that neutralizing VEGF caused partial recovery of barrier dysfunction, including decreased TER, enhanced sodium fluorescein flux, and lower ZO-1 expression induced by PAF. These results suggest that PAF-induced barrier dysfunction of RPE cells might be partly mediated by the upregulation of VEGF expression in RPE cells. The partial effectiveness of VEGF antibodies might indicate that factors other than VEGF might be involved in PAF-induced RPE barrier dysfunction. Our results contradicted the findings of Ghassemifar et al., who reported that VEGF slightly (approximately 10%) increased the TER of ARPE-19 cells and tightened RPE junctions[22]. The difference in results might be associated with the route or dose of VEGF administration.

ARPE-19 cells are unable to fully differentiate into RPE-like layers. For example, ARPE-19 cells have a low capacity to express critical proteins that regulate and maintain a barrier with strong tight junctions[15]. The use of ARPE-19 cells as a model of RPE function in vivo might be a limitation in the present study. Nevertheless, ARPE-19 cells remain valuable, as the changes appear to be quantitative rather than qualitative in terms of barrier function. Moreover, some characteristics of ARPE-19 cells, such as hypersensitivity to VEGF, loss of pigmentation, and weak tight junctions, somewhat resemble the characteristics of an aged eye or an eye with a pathologic condition[15].

## 5. Conclusion

In summary, the tight junctions of ARPE-19 cells are altered by PAF, and these effects are partly mediated by the upregulation of VEGF expression in RPE cells. Our results contribute to growing evidence supporting the role of PAF in choroidal neovascularization, and our findings suggest that PAF is a novel therapeutic target for increased permeability of RPE cells.

## Statements

### Acknowledgements

None.

### Statement of Ethics

This research was carried out on the ARPE-19 cell line in vitro and did not involve live human or animal subjects. The authors have no ethical conflicts to disclose.

### Authors’ contributions

Conceived and designed the experiments: HZ FZ LL ZLL. Performed the experiments: FZ HZ. Analyzed the data: LL HZ FZ ZLL. Contributed reagents/materials/analysis tools: FZ HZ. Wrote the paper: FZ LL HZ ZLL.

